# Paneth cells disruption and intestinal dysbiosis contribute to the development of Hirschsprung-associated enterocolitis in a benzalkonium chloride-induced Hirschsprung’s disease rat model

**DOI:** 10.1101/2023.08.19.553983

**Authors:** Iskandar Rahardjo Budianto, Kusmardi Kusmardi, Andi Muhammad Maulana, Somasundaram Arumugam, Rejina Afrin, Vivian Soetikno

## Abstract

**Background:** Hirschsprung-associated enterocolitis (HAEC) is a life-threatening complication of Hirschsprung’s disease (HSCR). This study investigated the role of Paneth cells (PCs) and gut microbiota in HAEC development.

**Methods:** Male Sprague-Dawley rats with HSCR were established by exposure of 0.1% (n = 30) benzalkonium chloride (BAC) to rectosigmoid serosa and sacrificed at 1-, 3-, 5-, 8-, and 12-weeks postintervention. The sham group was included and sacrificed on Week 12. Hematoxylin-Eosin staining was conducted to count the number of ganglionic cells and analyze the degree of enterocolitis. Intestinal barrier function was assessed for the ratio of anti-peripherin, occludin and acetylcholinesterase (AChE)/butyrylcholinesterase (BChE). PCs antimicrobial peptide (AMP) was evaluated by cryptdins, secretory Phospholipase A_2_, and lysozyme levels by qRT-PCR, respectively. 16S rRNA high throughput sequencing on faecal samples was used to analyze the changes in intestinal microbiota diversity in each group.

**Results:** Compared with sham groups, 0.1% BAC group rats had fewer ganglion cells after 1-week postintervention. Occludin and peripherin were decreased, and AChE/BChE ratio was increased, respectively. Sigmoid colon tissues from BAC-treated rats showed increased α-defensins positive PCs on Week 5 postintervention. Conversely, PCs-produced AMP tended to decrease from Week 5 to Week 12. Rats in the sham group demonstrated increased *Lactobacillus* and decreased *Bacteroides*, while rats in the 0.1% BAC exhibited reciprocal changes. Enterocolitis occurred from Week 1 postintervention onwards.

**Conclusion:** Disruption of PCs in the Week 5 postintervention and dysbiosis exacerbate the occurrence of HAEC. This research sheds new light on the cellular mechanisms of HAEC development.

## Introduction

Hirschsprung’s disease (HSCR) is a congenital malformation of the enteric nervous system characterized by the absence of ganglion cells in the distal colon with the resultant functional obstruction of the gut above the aganglionic segment and spastic contraction of the affected bowel [1]. One of the most severe complications of HSCR is Hirschsprung-associated enterocolitis (HAEC), which can occur both preoperatively and postoperatively [2,3].

HSCR animal models, piebald lethal and lethal spotted strains of mice, are the most frequently used animal models to study the pathophysiology of HSCR. Although the animal model of aganglionic genetics is very similar to the human HSCR condition, the aganglionic segment is concise and always located in the distal part of the rectum [4]. Moreover, those experimental animals die quickly from severe enterocolitis, making it difficult to study the complications of HSCR in those experimental animals [5]. Sato et al. have developed segmental aganglionosis through serosal topical application of benzalkonium chloride (BAC) to the colon and rectum of rats [6]. Myenteric plexus ablation and cholinergic nerve fibre hypertrophy resembling megacolon aganglionic [7] make this experimental animal model easy to use to study the pathophysiology and complications of HSCR.

Paneth cells, the specialized secretory epithelial cells in the mammalian small intestine, produce various secreted antimicrobial peptides (AMPs) that profoundly influence gut microbiota composition [8]. Dysfunction of Paneth cells compromises AMPs secretion leading to dysbiosis has been studied in various disease conditions, including inflammatory bowel disease (IBD) [9], acute necrotizing pancreatitis [10], necrotizing enterocolitis (NEC) [11], and chronic social defeat stress [12]. Pierre et al. demonstrated that *EdnrB*-null mice, transgenic mice that exhibit colonic aganglionosis, displayed dysbiosis, showed impaired mucosal defence, and decreased luminal secretory phospholipase A_2_ (sPLA_2_) before the development of HAEC [13]. A thorough examination of intestinal mucosal barrier dysfunction, Paneth cells and AMPs secretion, and dysbiosis in BAC-induced aganglionosis rats has not been undertaken. Therefore, in the present study, we investigated the role of Paneth cells and their AMPs secretion and changes in the gut microbiota associated with HAEC in BAC-induced aganglionosis rats by examining those changes at different times over 12 weeks.

## Materials and methods

### Animals

All animal testing protocols were approved by Animal Care and Use at Universitas Indonesia following the Universitas Indonesia Regulations of Animal Experimentation (ethical number: 471/UN2.F1/ETIK/PPM.00.02/2022). All surgeries were performed under 87 mg ketamine/kg of body weight and 13 mg xylazine/kg of body weight anaesthesia to minimize the suffering of rats.

Male Sprague-Dawley (SD) rats weighing 150 to 200 g were purchased from the Indonesian Food and Drug Authority, Indonesia. All rats were housed under conventional conditions maintained under a 12h light/dark cycle with water and food provided *ad libitum*. One cage contains one rat to facilitate the calculation of faeces’ weight every day.

Thirty-six SD rats were randomly divided into a sham-operated group (sham) (n = 6) and a 0.1% benzalkonium chloride (BAC) group (n = 30). The BAC group was further divided into five groups based on the time of termination, namely week 1 (W1), week 3 (W3), week 5 (W5), week 8 (W8), and week 12 (W12) post-BAC intervention. After induction of anaesthesia, laparotomy was performed via a 2 cm-long midline incision. After identification of the sigmoid colon, a 10×20 mm piece of paper gauze soaked in 0.1% of BAC was placed for 30 min around 1 cm of the sigmoid colon. Three drops of 0.1% BAC solution were given to the paper gauze every 5 min to prevent bowel dehydration. After 30 min of dripping BAC, the paper gauze was released, and the sigmoid colon was washed with 0.9% normal saline.

Then, the abdomen was closed, and all animals were housed individually with a standard diet and tap water ad libitum. The sham group rats underwent a sham operation, and the sigmoid colon was applied with 0.9% normal saline for 30 min.

### Postoperative evaluation

All rats were weighed once a week, while faeces were weighed daily. All rats were euthanasia by intraperitoneal injection with an overdose of ketamine/xylazine and were sacrificed by exsanguination under deep anaesthesia at week 1, week 3, week 5, week 8, and week 12 after BAC application. The isolated segment of the sigmoid colon was excised from each rat and divided into two parts, one part was promptly fixed in 10% neutral buffered formaldehyde solution for further histological examination, and the other part was washed with cold phosphate-buffered saline, snap-frozen in liquid nitrogen and stored at –80 °C for different experiments. Fresh faeces of rats in all groups were collected in sterile tubes and stored at –80 °C until further analysis.

### Histologic analysis

For light microscopy observation, the sigmoid colon segment from all rats fixed in 10% neutral buffered formaldehyde solution was dehydrated, embedded in paraffin and then cut into 5 μm sections. The sections were then stained with Hematoxylin-Eosin and analyzed by two pathologists blind to the study. The number of ganglionic cells per slice was counted and recorded. Teiltelbaum’s scoring was used to analyze the degree of enterocolitis histologically, namely Grade 0, normal intestinal mucosa; Grade 1, crypt dilation and mucin retention; Grade 2, cryptitis or crypt abscesses; Grade 3, multiple crypt abscesses; Grade 4, crypt hyperplasia and inflammatory cell infiltration, and Grade 5, crypt necrosis [14].

### Immunohistochemistry analysis for α-defensin

For immunohistochemistry (IHC), formalin-fixed, paraffin-embedded sigmoid colon tissue sections were used. After deparaffinization and hydration, the slides were washed in Tris-buffered saline (TBS; 10 mM/L Tris-HCl, 0.85% NaCl, pH 7.2). Endogenous peroxidase activity was quenched by incubating the slides with 0.3% H_2_O_2_ in methanol. After overnight incubation with the primary antibody, mouse monoclonal α-defensin 1 antibody (Novusbio, NBP2-75406, Centennial, USA), diluted 1:50, at 4 °C, the slides were washed in TBS followed by adding of HRP-conjugated recombinant anti-mouse antibody and incubated at room temperature for 45 min. The slides were then washed with TBS, incubated with diaminobenzidine tetrahydrochloride as a substrate, and then counterstained with hematoxylin. Semi-quantitative analysis of α-defensin 1 was carried out by counting the numbers of positive cells with strong expression.

### Immunofluorescence analysis for peripherin

Cryosections (5 μm) were cut using a microtome (Bright Instrument Company Ltd., Huntington, UK) housed within a cryostat at –25 °C. The sections were collected onto uncoated, precleaned glass slides fixed in 3.7% formaldehyde for one h. After rinsing in double distilled water for 3×30 sec, the slides were permeabilized in 0.5% Triton X-100 for 5 min. Subsequently, the slides were washed in phosphate-buffered saline (PBS, 137 mmol/L sodium chloride, 3 mmol/L potassium chloride, 8 mmol/L disodium hydrogen phosphate, and 3 mmol/L potassium dihydrogen phosphate, pH 7.4) for 3×5 min, and then the sections were incubated in anti-peripherin antibody (ab4666, Abcam, Cambridge, UK) for two h. Following a 3×5 min wash in PBS, a secondary Alexa Fluor fluorescent-conjugated antibody was applied for 30 min. Secondary antibody controls were included as negative controls. Images were taken with a fluorescence microscope (Olympus, BX43, Japan).

### Gene expression analysis for α-defensin, lysozyme, sPLA2, and interleukin (IL)-1β

Total RNA was extracted from sigmoid colon tissue using a High Pure RNA Isolation kit (Roche Applied Science, Penzberg, Germany) according to the manufacturer’s instructions. Nanodrop 1000 Spectrophotometer (Thermo Scientific) was used to measure RNA concentration at a wavelength of 260 nm. 1 μg of RNA, and the Transcriptor First Strand cDNA Synthesis kit (Roche Applied Science) were used to synthesize complementary DNA according to manufacturer’s instructions. Quantitative Real-Time PCR (qRT-PCR) was done using the LightCycler® 480 Instrument (Roche Applied Science) with FastStart Essential DNA Green Master Mix (Roche Life Science). All reactions were performed similarly: 95 °C for 10 sec, followed by 45 cycles of 95 °C for 15 sec and 60 °C for 1 min. Relative gene expression was calculated using the 2^-ΔΔCT^ method, with β-actin as the housekeeping gene. The following primer pairs were used: β-actin: Forward: 5’-CTGGTCGTACCACAGGCATT-3’ Reverse: 5’-CTCTTTGATGTCACGCACGA-3’; α-defensin (cryptdins-1): Forward: 5’-CCG AGA GTG CTT CCT AAA CTA C-3’ Reverse: 5’-AAA GTC TCA GGT GGG ATG TTA G-3’; lysozyme: Forward: 5’-GAA TGG GAT GTC TGG CTA CTA TG-3’ Reverse: 5’-GTC TCC AGG GTT GTA GTT TCT G-3’; sPLA2: Forward: 5’-ACT CAT GAC CAC TGC TAC AAT C-3’ Reverse: 5’-GTA TGA GTA CGT GTT GGT GTA GG-3’; IL-1β: Forward: 5’-CCA GGA TGA GGA CCC AAG CA-3’ Reverse: 5’-TCC CGA CCA TTG CTG TTT CC-3’

### Western blotting analysis for AChE, BChE, and occludin antibodies

The frozen sigmoid colonic tissues were weighed and homogenized in an ice-cold Tris buffer (50 mM Tris-HCl, pH 7.4, 200 mM NaCl, 20 mM NaF, 1 mM Na_3_VO_4_, 1 mM 2-mercaptoethanol, 0.01 mg/mL leupeptin, 0.01 mg/mL aprotinin). After that, homogenates were centrifuged (3000 x *g*, 10 min, 4 °C) and the supernatant were collected and stored at –80 °C. The total protein concentration in each sample was measured by the bicinchoninic acid method. Equal amounts of protein extract (50 μg) were separated by sodium dodecyl sulfate-polyacrylamide gel electrophoresis (Bio-Rad, CA, USA) and transferred electrophoretically to nitrocellulose membranes. Then, the membranes were blocked with 5% skim milk in Tris-buffered saline Tween 20 (20 mM Tris, pH 7.6, 137 mM NaCl and 0.1% Tween 20). Subsequently, membranes were incubated overnight at 4 °C with agitation with primary antibodies against acetylcholinesterase (68 kDa, 1:500, ab97299; Abcam, Cambridge, UK), butyrylcholinesterase (75 kDa, 1:500, AF9024; R&D systems, USA), and occludin (60 kDa, 1:1000, sc-133256; Santa Cruz, CA, USA). Membranes were washed three times in TBS-T for 15 min and incubated with horseradish peroxidase-conjugated goat anti-rabbit IgG secondary antibody (1:5000, Santa Cruz) or rabbit anti-mouse IgG secondary antibody (1:5000, Santa Cruz) for one h at room temperature. After a final wash, signals were detected using a chemiluminescence system (ECL western blotting substrate, Life Technologies). The expression of the above antibodies was normalized by glyceraldehyde 3-phosphate dehydrogenase (GAPDH) protein expression in the same sample. At last, membranes were scanned and analyzed using ImageJ software.

### Microbiota analysis

Bacterial DNA was extracted from 200 mg rat faeces per sample with the ZymoBIOMICS^TM^ DNA Miniprep Kit (catalogue #D4300) according to the manufacturer’s protocols. Bacterial DNA was amplified with 341F (CCTAYGGGRBGCASCAG) and 806R (GGACTACNNGGGTATCTAAT) primers, including the V3-V4 region of the bacterial 16S rRNA gene. The PCR products with the proper size were selected by 2% agarose gel electrophoresis. The exact amount of PCR products from each sample was pooled, end-repaired, A-tailed, and ligated with illumine adapters. Libraries were sequenced on a paired-end Illumina platform to generate 250bp paired-end raw reads. The library was checked with Qubit and real-time PCR for quantification and a bioanalyzer for size distribution detection. Quantified libraries were pooled and sequenced on the Illumina platform according to the required effective library concentration and data amount.

### Statistical analysis

Data were analyzed using Prism GraphPad version 9.5.1 software. Data were reported as the mean ± standard error of the mean (SEM). Statistical significance was determined using the one-way ANOVA followed by Tukey’s posthoc for multiple comparisons. Differences among groups were considered significant at p < 0.05.

## Results

### Body weight, faeces weight, and abdominal circumference changes among groups

All rats survived until the end of the study. At the end of the study, the body weight in the BAC-treated groups was significantly lower than in the sham group (Fig. 1A). Faeces weight in the BAC-treated group was also lower than in the sham group. However, the difference was significant only in the Week 3 BAC-treated group (Fig. 1B). BAC administration causes manifestations of hypoganglionosis, namely abdominal distention, which increases according to the time of termination, as seen in the increase in abdominal circumference (Fig.1C).

**Figure 1.**
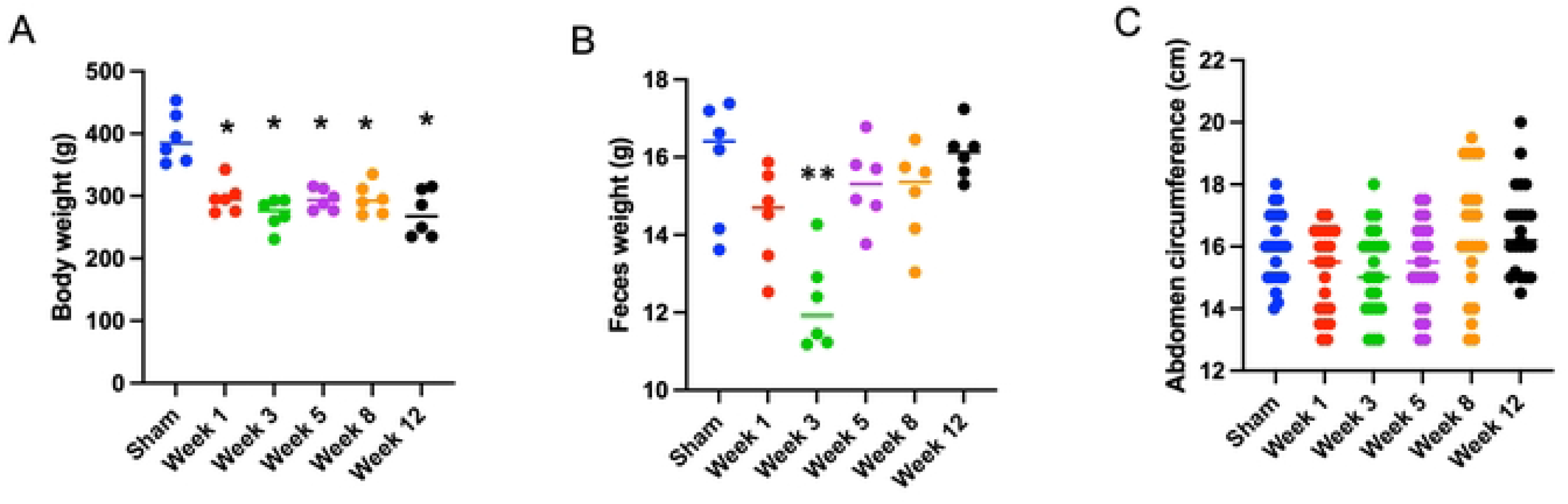
Body weight, faeces weight, and abdomen circumference in the sham and BAC-treated groups terminated on Week 1 – Week 12 postintervention. (A) Distribution of body weight in each group of rats. The body weight of BAC-treated rats was significantly lower than the sham group (each, *n* = 6). (B) Distribution of faeces weight in each group of rats. There was no difference in faeces weight in BAC-treated rats as compared to the sham group, though faeces weight in the Week 3 of BAC-treated rats decreased significantly as compared to the sham group (p < 0.01) (each, *n* = 6). (C) Distribution of abdomen circumference in each group of rats. There was no difference in abdomen circumference in BAC-treated rats compared to the sham group.

### Ganglionic cells number among groups

As shown in Fig. 2A, the number of ganglion cells per ganglia was reduced in the BAC-treated group compared to the sham group. This was in accordance with the quantitative analysis in Fig. 2B, which showed that the mean number of ganglion cells per ganglia decreases with the time of termination. Peripherin, a specific marker for enteric neuronal cells (ganglia), showed ganglia in all groups. The expression of ganglia appeared to be reduced in the BAC-treated group according to time of termination compared to the sham group (Fig. 2C).

**Figure 2.**
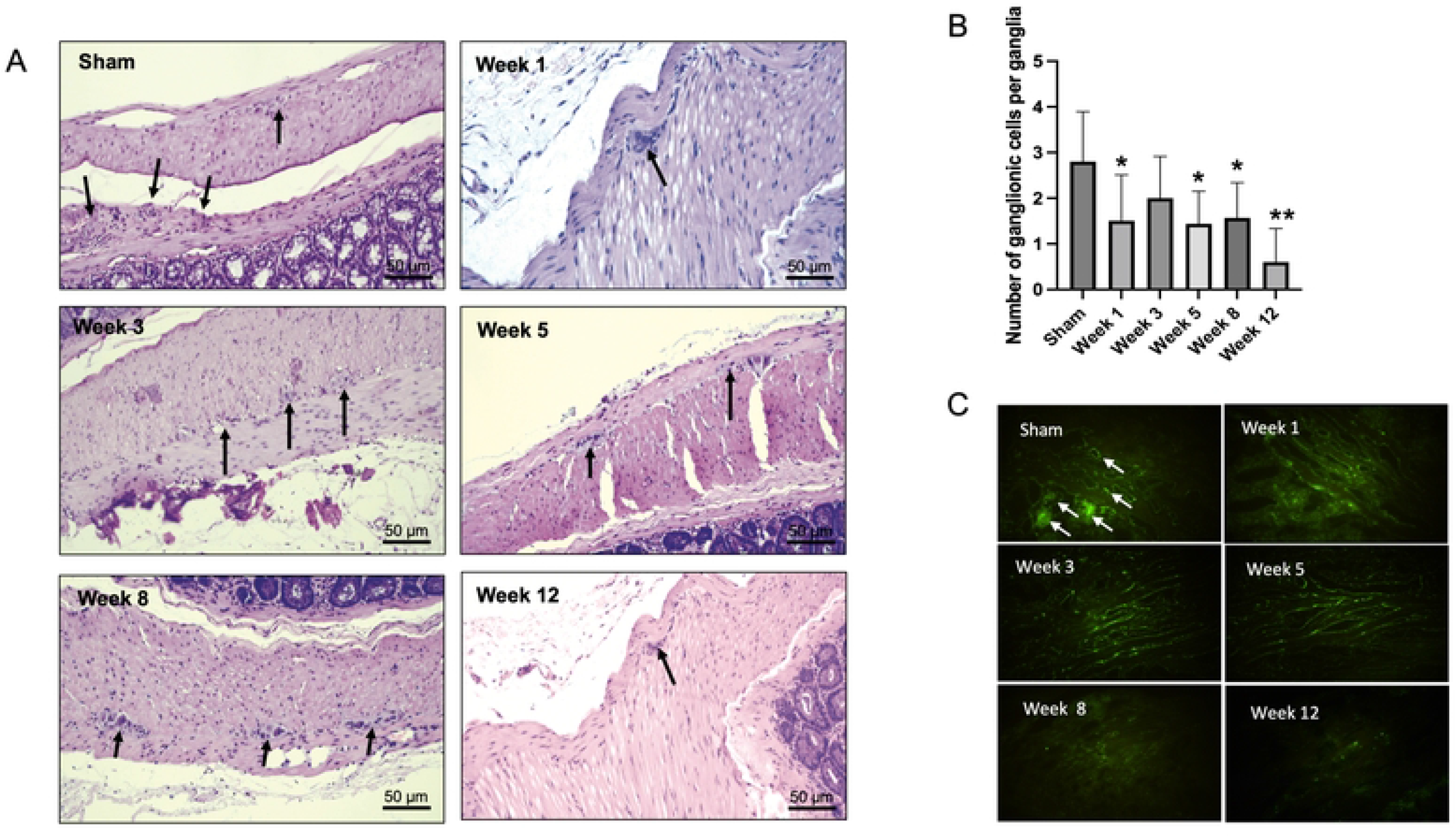
Hematoxylin and eosin (H&E) staining of the rectosigmoid colon in the sham group and BAC-treated group terminated on Week 1 – Week 12 postintervention. (A) H&E staining showing ganglionic cells in a 1-cm/5-μm slice of the myenteric plexus. **(B)** Compared to the sham group, a decreased mean number of ganglionic cells occurred at Week 1 to Week 12 post-denervation. **(C)** Immunofluorescence staining of peripherin, a neuronal marker, showed ganglionic cells in the sham and BAC-treated groups terminated on Week 1 – Week 12. Magnification x40 for A and C (Scale bar = 50 μm). The data of B was shown as the mean ± SE of six independent experiments in each group. A one-way ANOVA test followed by Tukey’s test was used to compare the data. *p<0.05 vs sham; **p<0.01 vs sham.

### Changes in intestinal barrier permeability among groups

We assessed the AChE/BChE ratio to analyze mucosal cholinergic innervation. As shown in Fig. 3A, the ratio of AChE/BChE was higher in the BAC-treated groups than the sham group, mainly at Week 5 postintervention. An increase in the AChE/BChE ratio indicates extrinsic nerve fibre hypertrophy in the aganglionic segment. Next, to assess intestinal permeability, we analyzed the protein expression of the tight junction, occludin, by Western blot. Expression of occludin decreased in the BAC-treated group according to the time of termination as compared to the sham group (Fig. 3B). To analyze the presence of an inflammatory process due to the loss of intestinal tight junction barrier function, we measured the gene expression of IL-1β, which was increased in the BAC-treated group, mainly at Week 8 postintervention (3-fold) than the sham group (Fig. 3C).

**Figure 3.**
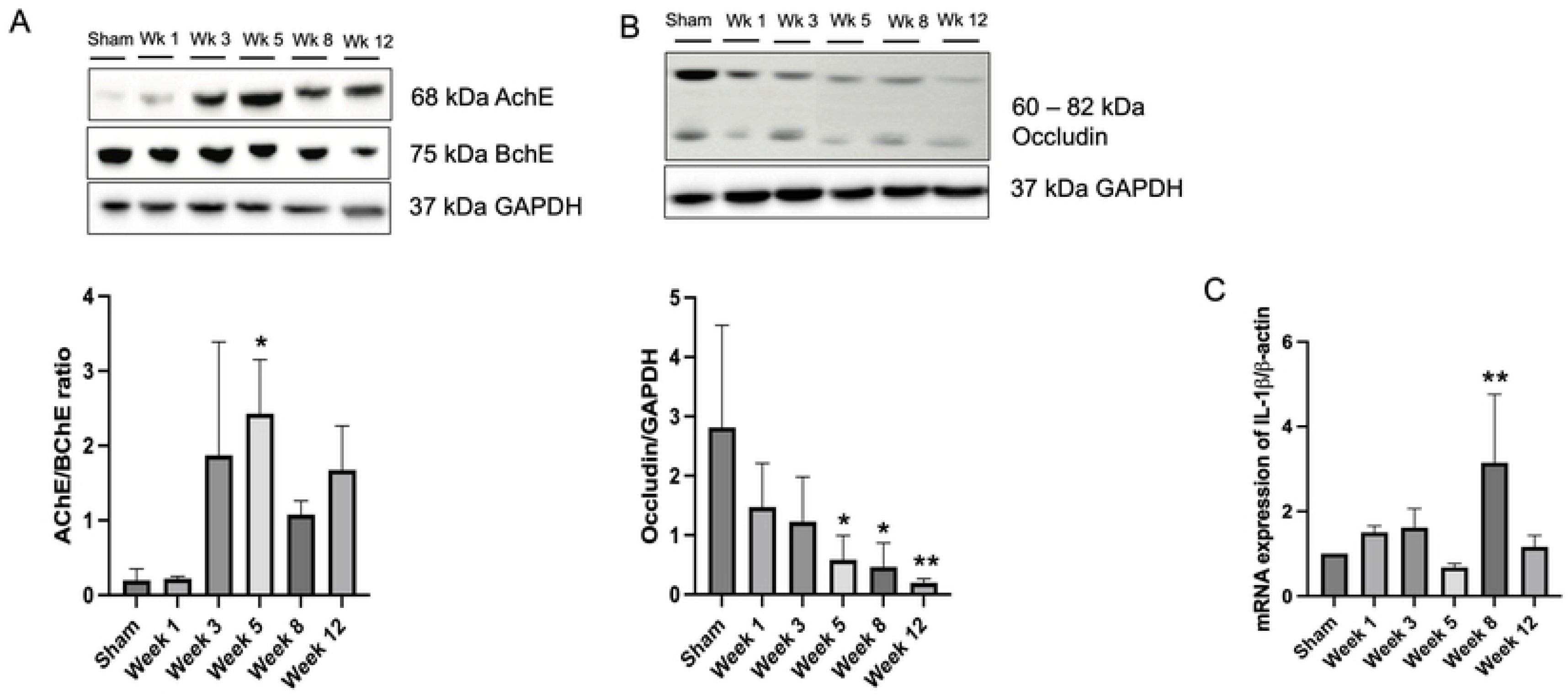
Benzalkonium chloride-induced aganglionosis results in hypertrophy of cholinergic nerve fibre, loss of intestinal tight junctions, and inflammation. (A) Western blotting revealed an increased AChE/BChE ratio in the rectosigmoid colon of BAC-treated groups (n = 6) compared to the sham group (n = 6) according to the time of termination. **(B)** Western blotting revealed decreased occludin tight junction in the rectosigmoid colon of BAC-treated groups (n = 6) compared to the sham group (n = 6) according to the time of termination. **(C)** mRNA expression of IL-1β on rectosigmoid colon tissue between groups. Results are presented as mean ± SE. A one-way ANOVA test followed by Tukey’s test was used to compare the data. *p<0.05 vs sham; **p<0.01 vs sham.

### Modulation of Paneth cells antimicrobial peptides in BAC-treated rats

We conducted immunohistochemical analysis of α-defensin to detect Paneth cells in the sigmoid colon tissue and found that the α-defensin positive Paneth cells were almost absent in the sham group, whereas in the BAC-treated groups, the number of α-defensin positive Paneth cells continued to increase according to the time of termination and reached a peak at Week 5 postintervention, which proved that Paneth cells metaplasia occurred in the sigmoid colon tissue of HSCR rat model (Fig. 4A & 4B).

**Figure 4.**
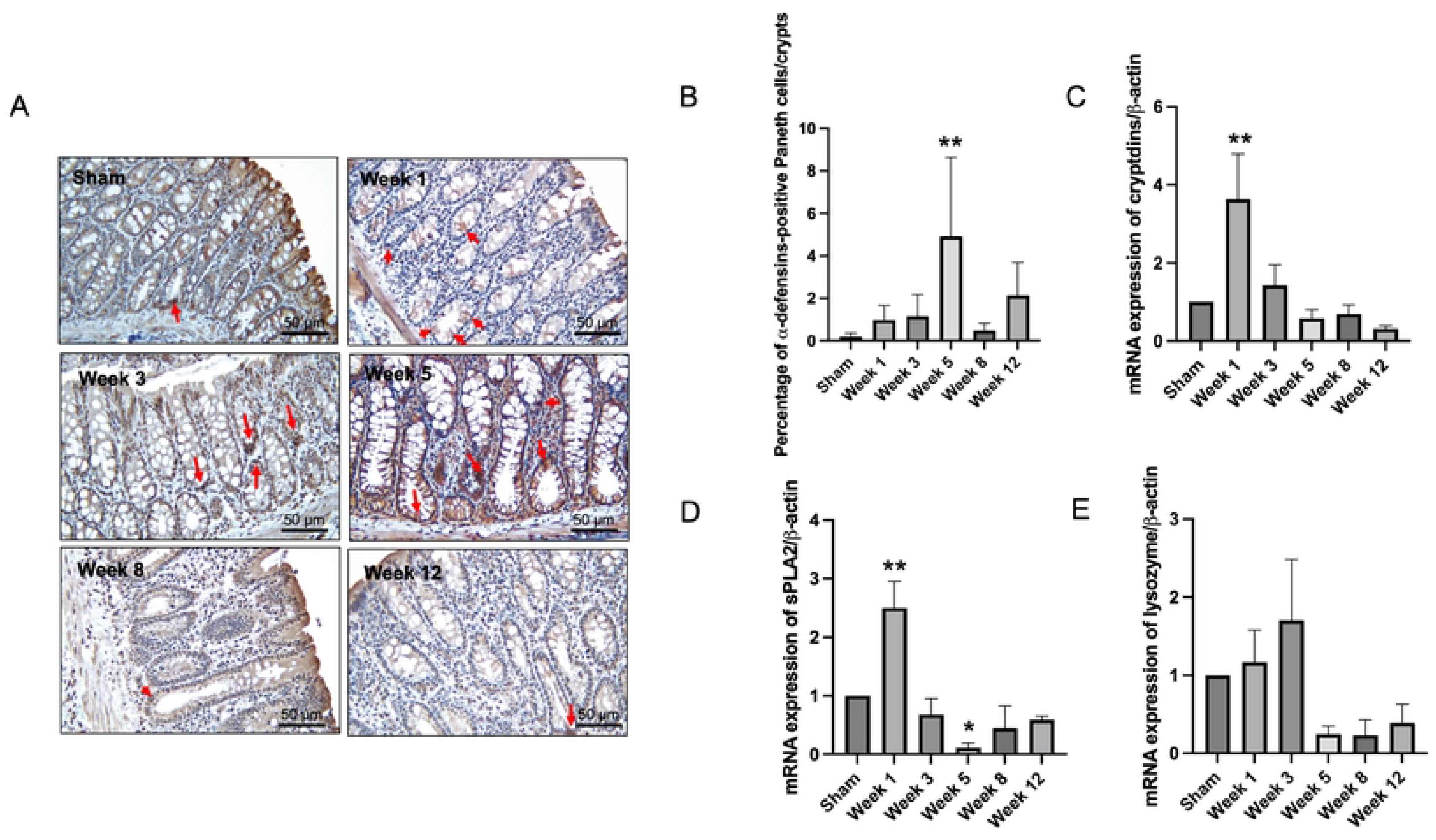
Paneth cells metaplasia in the rectosigmoid colon and mRNA expressions of Paneth cells-secreted antimicrobial peptides (A) Immunohistochemistry for α-defensins-positive Paneth cells in rectosigmoid tissue from the sham group and BAC-treated groups. Red arrows indicate α-defensins-positive Paneth cells. Magnification x40 (Scale bar = 50 μm). **(B)** Quantitative analysis of α-defensins-positive Paneth cells. The sham group showed few Paneth cells, while the BAC-treated group showed increased Paneth cells, mainly in Week 5 in the rectosigmoid colon. The data are shown as the mean ± SE of six independent experiments in each group. A one-way ANOVA test followed by Tukey’s test was used to compare the data. **(C)** mRNA expression of cryptdins, sPLA_2_ **(D)**, and lysozyme **(E)** showed a cycle, increased at Week 1, indicating Paneth cells turnover, followed by a decrease until Week 12, which indicates Paneth cells dysfunction. The data are shown as the mean ± SE of six independent experiments in each group. A one-way ANOVA test followed by Tukey’s test was used to compare the data. **p<0.01 vs sham.

Next, we analyzed the antimicrobial peptide secreted by Paneth cells, namely α-defensin (cryptdin), sPLA2, and lysozyme by qRT-PCR. As shown in Fig. 4C-E, the gene expression levels of α-defensin (cryptdin), sPLA2, and lysozyme started to increase at Week 1 postintervention and then decreased until the end of the study.

### Composition of luminal microbiota among groups

The composition of luminal microbiota in the rectosigmoid of the BAC-treated and sham groups was assessed by 16SrDNA sequencing each of 4 animals. Hierarchical clustering analysis was performed at each taxonomic level from phylum through genus at the time of termination points. At the phylum level, *Firmicutes* and *Bacteroidetes* were the two major bacterial phyla in the gastrointestinal tract of both the sham and the BAC-treated groups (Fig. 5A). As shown in Fig. 5B, the *Firmicutes/Bacteroidetes* (F/B) ratio decreased in the BAC-treated group compared to the sham group. The F/B ratio appeared to be lower according to the time of termination. To further understand the phyla contributing to these grouping, the top three phyla found at each time point of termination were analyzed between groups (Fig. 5C). The sham group demonstrated decreased representation of *Bacteroidetes* and *Proteobacteria*, with increased *Firmicutes*. Conversely, in the BAC-treated group, *Bacteroidetes* and *Proteobacteria* increased, and *Firmicutes* decreased, mainly at Week 8 – Week 12 postintervention, though there were no significant differences between groups (Fig. 5C).

**Figure 5.**
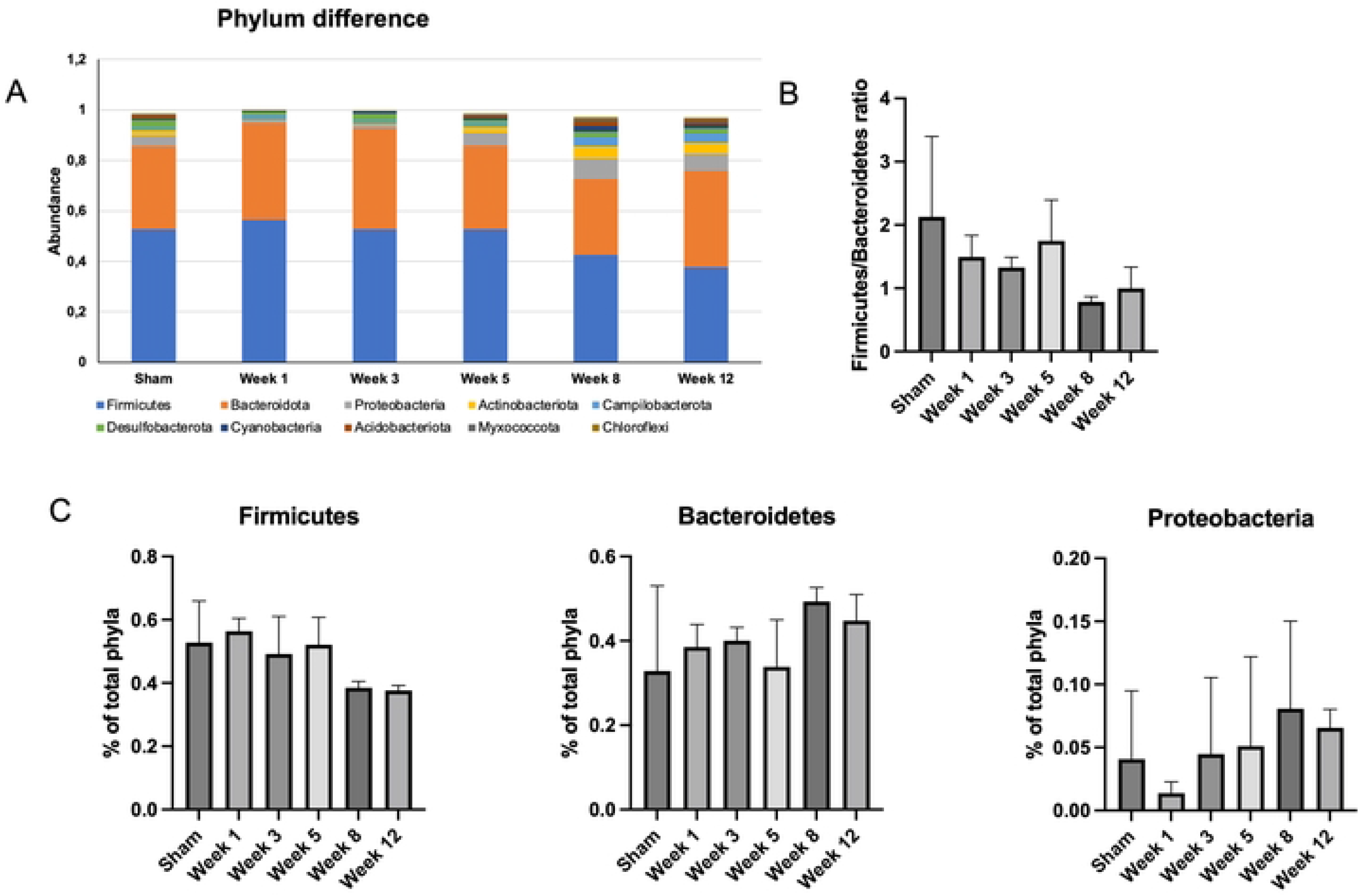
Changes in gut microbiota diversity and structure in HSCR at the phylum taxonomic. (A) Diversity of gut microbiota at phylum levels. **(B)** *Firmicutes/Bacteroidetes* ratio among groups showed a significantly decreased F/B ratio at Week 8 – 12 postintervention **(C)** Relative abundance of different phyla among groups. *Bacteroidetes* and *Proteobacteria* increased in the BAC-treated groups, while *Firmicutes* increased in the sham group at Week 8 – 12 postintervention. The data are shown as the mean ± SE of four independent experiments in each group. A one-way ANOVA test followed by Tukey’s test was used to compare the data.

At the genus level, the top ten genera in each group are shown in Fig. 6A. Prevotella is the most common genus found in the sham and BAC-treated groups. *Lactobacillus*, a species considered to provide intestinal mucosal protection, was higher in the sham group and decreased in the BAC-treated group. In contrast, *Bacteroides* increased significantly at Week 8 postintervention compared to the other groups (Fig. 6B).

**Figure 6.**
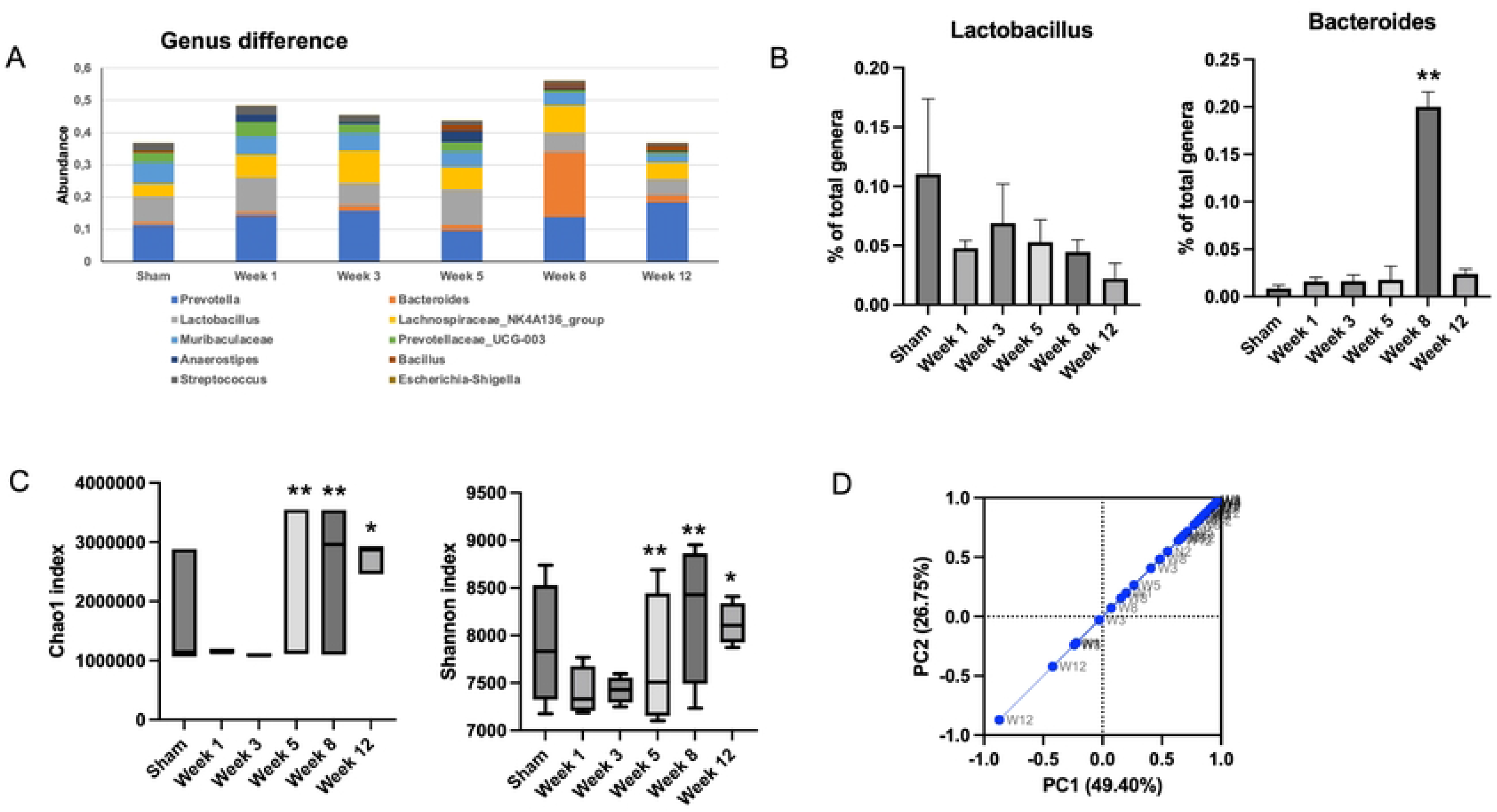
Changes in gut microbiota diversity in HSCR at the genus taxonomic. (A) Diversity of gut microbiota at genus levels. **(B)** Relative abundance of *Lactobacillus* and *Bacteroides* among groups. *Lactobacillus* increased in the sham group, while *Bacteroides* increased in the BAC-treated groups at Week 8 postintervention. The data are shown as the mean ± SE of four independent experiments in each group. A one-way ANOVA test followed by Tukey’s test was used to compare the data. **(C)** Difference of alpha diversity indices between groups. The richness and diversity significantly differed between early (Week 1 and Week 3) and late weeks (Week 5 – Week 12). ^$^p<0.05 vs W301; ^#^p<0.05 vs W101; *p<0.05 vs W802. **(D)** Difference of beta diversity indices between groups. The composition of microbial communities differed between sham and BAC-treated groups, mainly at Week 12 postintervention.

### Microbiota diversity and richness

Richness (Chao1index) and diversity (Shannon index) were similar between the sham group and BAC-treated group (Fig. 6C). However, the richness and diversity of microbial communities were significantly different between early weeks (Week 1 and Week 3) and late weeks (Week 5 to Week 12) postintervention of BAC. As shown in Fig. 6D, PCA described the β diversity, revealing a separation between the sham group and the BAC-treated group at Week 8 – Week 12. PCA showed that PC1 and PC2 accounted for 49.40% and 26.75% variance, respectively, and PC1 clearly separated the samples in the BAC-treated group in the early and late weeks. This result implied that there were differences in the structure of the gut microbiota in early and advanced HSCR.

### Degree of enterocolitis among groups

Degree of enterocolitis grade 1 – grade 5 was found in the BAC-treated group (Fig. 7Ai-vi). Grade 1 degree of enterocolitis began to occur in Week 1 postintervention, and the degree of enterocolitis increased according to the time of termination (Fig. 7B).

**Figure 7.**
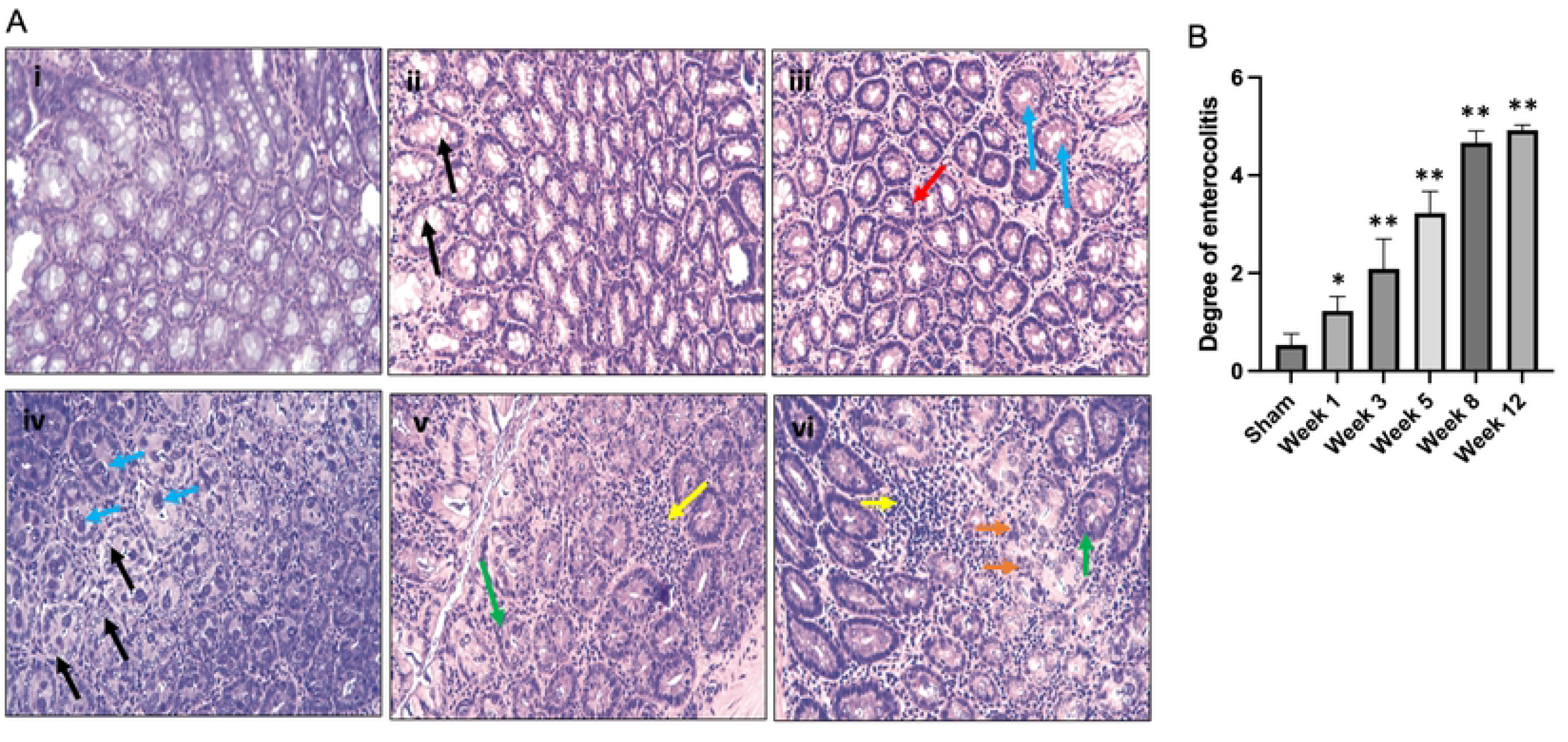
(A) Degree of Hirschsprung-associated enterocolitis. [i] Grade 0, normal mucosa. [ii] Grade 1, crypt dilation with mucin retention (black arrows). [iii] Grade 2, cryptitis (red arrow) and crypt abscesses (blue arrows). [iv] Grade 3, crypt abscesses (blue arrows) and crypt dilatation with mucin retention (black arrows). [v] Grade 4, inflammatory cell infiltration (yellow arrow) and crypt hyperplasia (green arrow). [vi] Grade 5, inflammatory cell infiltration (yellow arrow), crypt necrosis (orange arrows), and crypt hyperplasia (red arrow). **(B) Degree of Hirschsprung-associated enterocolitis between groups.** The data are shown as the mean ± SE of six independent experiments in each group. A one-way ANOVA test followed by Tukey’s test was used to compare the data.

## Discussion

In the present study, we hypothesized that Paneth cells and dysbiosis have an important role in the development of HAEC in the BAC-induced aganglionosis HSCR rat model. To induce the HSCR model in rats, we used a BAC concentration of 0.1%. We found that the application of 0.1% BAC caused a decrease in the number of nerve ganglion cells and disturbed intestinal permeability in an exposure time-dependent manner. We also showed that rats that received BAC experienced abdominal distension and loose faeces, especially in late weeks, and decreased body weight. In addition, we also identified that the number of Paneth cells began to increase in the first week, then the number greatly increased at Week 5, followed by a decrease until Week 12.

Serosal application of BAC with various concentrations has been widely used to induce HSCR in experimental animals [6,15–17]. In the present study, we used 0.1% BAC concentrations to determine myenteric denervation and aganglionosis. Our procedure results in robust aganglionosis at 0.1% BAC. In addition, histopathology analysis showed neuronal ganglion cell ablation beginning in the first week and persisting until the 12^th^ week after denervation. Unlike our research, previous studies have reported spontaneous regeneration of enteric ganglion cells in the first two months after myenteric denervation [18,19].

In the present study, we observed an increase in the AChE/BChE ratio in the group that received 0.1% of BAC compared to the sham group, which indicated parasympathetic nerve fibre hypertrophy. Prior studies have shown higher AChE activity in HSCR, in which determination of the AChE/BChE ratio may have discriminatory diagnostic value; the AChE/BChE ratio was markedly decreased in cholinergic nerve fibres when ganglion cells were present [20–22]. Moreover, Keck et al. have demonstrated an increased AChE activity in the rectosigmoid of HSCR patients [23].

Enterocyte tight junction proteins play a critical role in maintaining intestinal homeostasis and act as a barrier yet absorb substances paracellularly. Some enterocolitis conditions cause impaired epithelial tight junction barrier function [24,25]. Occludin is one of the tight junctions necessary to maintain intestinal integrity and as a transmembrane signalling protein [26]. Our data showed that occludin was downregulated in the rectosigmoid colon of the BAC-treated group, mainly in the Week 5 postintervention. Consistent with our results, Arnaud et al. demonstrated that tight junction protein zonula occludens-1 expression was significantly decreased in piglets with hypo ganglionic sigmoid [16]. It has been shown that reduced occludin expression in both epithelial and endothelial cells was related to increased intestinal barrier permeability [27,28]. Dysregulated epithelial tight junction protein and the consequent increase of intestinal permeability and gut dysbiosis correlate with the development and progression of pathological conditions such as HSCR and necrotizing enterocolitis (NEC) [29,30]. Previous studies have shown that proinflammatory cytokines, including IL-1β were markedly elevated in IBD and NEC [31–33]. In line with the results of previous studies linking intestinal epithelial tight junction barrier loss with inflammatory processes, we also prove that the proinflammatory cytokine IL-1β increased in the BAC-treated group, is probably due to the loss of intestinal epithelial tight junction protein.

Paneth cells play a key role in the intestinal innate host defence in which they secrete antimicrobial peptides (AMPs) in response to bacteria or their antigens [34]. Previous studies have demonstrated that Paneth cells dysfunction and Paneth cells metaplasia and hyperplasia occurred in NEC and IBD in association with inflammatory processes in both diseases [11,35]. We have recently reported that Paneth cells were hyperplastic and metaplastic in the sigmoid colon of HSCR rats [36]. In the present study, we found Paneth cells metaplasia in the rectosigmoid colon of the BAC-treated group, and the number greatly increased at Week 5 postintervention, followed by a decrease until Week 12 postintervention. In contrast, very few Paneth cells numbers could be identified in the sham group.

Furthermore, we analyzed the alterations in gene expression of lysozyme, sPLA2, and cryptdins, the major Paneth cells AMPs. Our results showed that all Paneth cells AMPs’ gene expression increased at Week 1 postintervention and decreased until Week 12 postintervention. The increase in the number of α-defensins-positive Paneth cells in Week 1 corresponded to the rise of Paneth cell number. However, we found that Paneth cells became dysfunctional at Week 5 postintervention, whereby Paneth cell synthesis occurred without the ability to secrete AMPs. In previous research, Imamura et al. observed a decrease in luminal sIgA along the resected aganglionic colon despite an increase in sIgA-containing plasma cells in HSCR patients with HAEC compared to patients without HAEC [37]. Impaired Paneth cell function also occurs in experimental animals with intestinal ischemia/reperfusion [38] and acute kidney injury, in which acute loss of renal function causes Paneth cells-induced release of IL-17A and intestinal damage [39]. Recent investigations have identified numerous diseases that can disrupt normal Paneth cell function, resulting in compromised AMPs secretion and consequent dysbiosis [40–42].

The intestinal barrier mediates crosstalk between commensal intestinal microbes and host immunity and is the first line of defence against pathogenic antigens and potentially harmful microorganisms [43]. Thus, the impaired intestinal barrier may lead to pathogen invasion and mucosal dysbiosis, which ultimately trigger pathologic conditions, including IBD and HAEC [13,44]. In our study, the microbial composition of faeces from BAC-treated groups was shifted toward opportunistic pathogens such as *Proteobacteria*. A shift induced by HAEC in the gut microbiota structure was reflected by PCA, in which the composition of microbial communities differed between the sham group and BAC-treated groups, mainly in the late week of HAEC. However, our data showed no microbial abundance and diversity differences between the sham and BAC-treated groups in the early week. In contrast, these differences appeared in the late week, as indicated by α-diversity.

Similarly, Pierre et al. proved that Chao-1 and Shannon’s index were similar between the control and EdnrB-null mice groups at an early stage [13]. Expansion of *Proteobacteria* is an important finding in HSCR patients, especially with recurrent HAEC. *Proteobacteria* have been reported to significantly impact the pathogenesis of HAEC and IBD [45–47]. Moreover, it has been known that the Firmicutes/Bacteroidetes (F/B) ratio was associated with the maintenance of intestinal homeostasis, and changes in this ratio can result in various pathologies, such as increases in the abundance of *Firmicutes* or *Bacteroidetes* species lead to obesity and IBD, respectively [48,49]. In our present study, we found that the F/B ratio tended to decrease in the BAC-treated group compared to the sham group. We also demonstrated that enterocolitis occurred at Week 1 postintervention, and the degree of enterocolitis continued to increase according to the time of termination. We believe this was due to Paneth cells dysfunction and dysbiosis that exacerbate HAEC.

We are perfectly conscious that one limitation of our study is that there is no sham group at each time of termination. Therefore, the differences in each parameter with the group that received BAC treatment cannot be ascertained.

Together our results suggest that the impaired intestinal mucosal barrier and disruption of Paneth cell functions in maintaining the intestinal tract’s stability are associated with dysbiosis in HAEC. These findings further confirm the involvement of Paneth cells and gut microbes in the multifactorial aetiology of HSCR and its complication, HAEC, which may facilitate the development of management for prevention and treatment.

## Acknowledgements

We thank Chyswyta, Annisa, Anggari Kirana Dewi, and Mariatul Qiftia for helping with technical assistance.

